# STRAND: Structure Refinement of RNA-Protein Complexes via Diffusion

**DOI:** 10.1101/2025.06.30.662415

**Authors:** Mohsen Al-zeqri, Jörg K.H. Franke, Frederic Runge

**Affiliations:** Department of Computer Science, University of Freiburg, Germany; ELLIS Institute Tü bingen, Germany; LAION

## Abstract

RNA-protein interactions play crucial roles in cellular processes, from gene regulation to viral replication. While recent advances in structure prediction have revolutionized our ability to model macromolecular complexes, achieving accurate predictions of RNA-protein binding poses remains challenging. In this work, we present *STRAND*, a diffusion-based model for monomeric RNA-protein complex refinement that builds upon the success of *DiffDock-PP* in protein-protein docking. Unlike traditional docking, we develop STRAND as a modular extension to existing RNA-Protein complex prediction tools to improve their backbone predictions. We study the effect of different transformations by training models to learn either translation, rotation, torsion, or combinations of these during the diffusion process and initialize the backward process with a complex prediction at test time. Our experiments with *AlphaFold 3* and *ProRNA3D-single* reveal that STRAND can improve the backbones of a large fraction of RNA-protein complex predictions.

## 1. Introduction

RNA-protein interactions are fundamental to numerous cellular processes, including transcriptional regulation, splicing, and protein synthesis (Glisovic et al., 2008). Understanding these interactions at the structural level is crucial for deciphering molecular mechanisms and developing therapeutic interventions. While experimental techniques like X-ray crystallography (Smyth & Martin, 2000), nuclear magnetic resonance spectroscopy (Hu et al., 2021), or single-particle electron microscopy (cryo-EM) (Renaud et al., 2018) provide valuable structural insights, they are often time-consuming and resource-intensive.

Recent breakthroughs in deep learning have revolutionized structural biology, particularly with the advent of AlphaFold 3, which enables predictions of RNA-protein complexes alongside other biological macromolecules (Abramson et al., 2024). However, while deep learning methods have driven progress in RNA secondary structure prediction (Sato et al., 2021; Franke et al., 2024) and secondary structure-(Runge et al., 2024; Patil et al., 2024) as well as 3D-based (Tan et al., 2024; Joshi et al., 2025) RNA design, RNA 3D structure prediction remains challenging (Bernard et al., 2025). This challenge is compounded by the inherent flexibility of RNA molecules (Hagerman, 1997; Franke et al., 2022) and the diverse nature of RNA-protein binding interfaces (Re et al., 2014).

The recent success of diffusion models in protein-protein docking, particularly DiffDock-PP (Ketata et al., 2023), has demonstrated the potential of these approaches in modeling molecular interactions. However, these methods are specifically designed for protein-protein interfaces and do not account for the distinct characteristics of RNA molecules.

In this work, we present STRAND (**ST**ructure **R**efinement of RN**A**-protei**N** complexes via **D**iffusion), a novel diffusion-based model that extends the capabilities of DiffDock-PP to monomeric RNA-protein complexes. Our approach leverages an RNA foundation model, RNA-FM (Chen et al., 2022), to generate RNA sequence embeddings that capture the unique properties of RNA molecules. We then combine these RNA-specific features with the powerful diffusion framework of DiffDock-PP. We train different models, applying noise either on the translation level, the rotation level, to torsion angles, or combinations of these across thousands of experimentally validated RNA-Protein complexes from the Protein Data Bank (PDB) (Burley et al., 2017). Then, at test time, we use RNA-Protein complex predictions from AlphaFold 3 (Abramson et al., 2024) or ProRNA3D-single (Roche et al., 2024) as the initial complex structure for the denoising process to refine the initial predictions. With this approach, STRAND is capable of improving many initial predictions, as indicated by improved complex RMSD (cRMSD) scores. STRAND thus represents a novel approach to RNA-protein complex refinement that leverages recent advancements in 3D RNA-protein modeling, and we propose to use STRAND as a module on top of existing RNA-protein complex prediction tools to enhance their predictions.

Our main contributions can be summarized as follows:

We propose STRAND, a novel RNA-protein refinement strategy that leverages the new modeling capabilities of current state-of-the-art deep learning based RNA-protein complex prediction approaches by directly improving their predictions. STRAND represents a modular extension to existing RNA-Protein complex prediction approaches.

- We extend DiffDock-PP to the prediction of RNA-Protein complexes using the embeddings of an RNA foundation model, RNA-FM (Section 3).
- Our experiments reveal improvements in complex structure prediction compared to the initial predictions of AlphaFold 3 and ProRNA3D-single across a diverse set of RNA-protein complexes from the PDB (Section 4).
- We show predictions of STRAND trained with translational noise only that exemplify structural improvements together with initial predictions of AlphaFold 3, superimposed on the original PDB structure in Figure 1.

**Figure 1.**
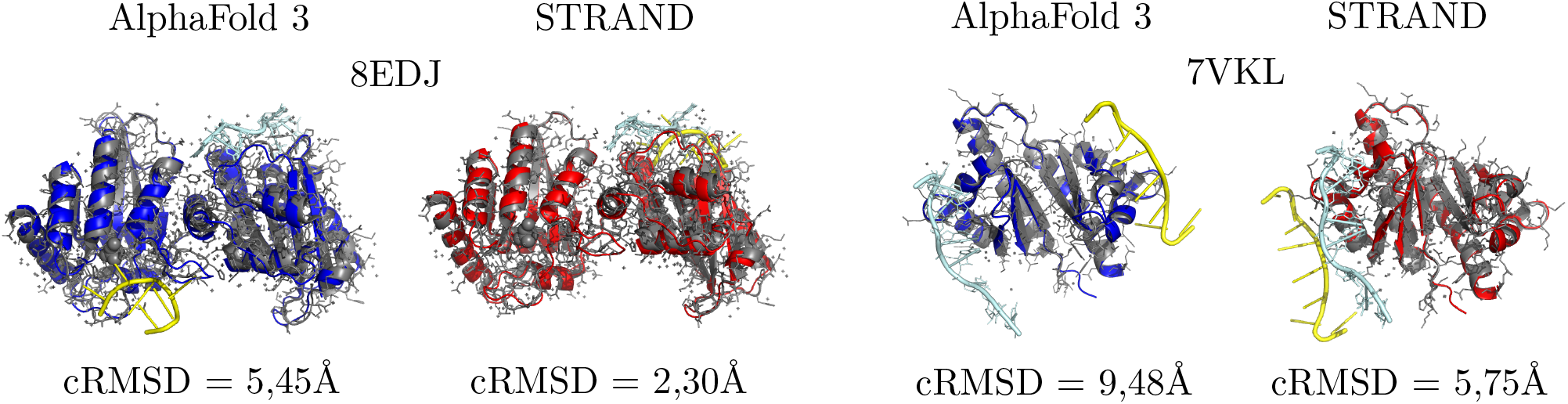
Example structure refinements for two AlphaFold 3 predictions for the PDB Ids 8EDJ and 7VKL using STRAND trained only on translational noise. We show the predictions of AlphaFold 3 (blue) and STRAND*_tr_*(red) superimposed on the true PDB structure of 8EDJ (left) and 7VKL (right). The true protein structure is shown in gray, the RNA in light blue. The RNA of the prediction is shown in yellow.

## 2. Related Work

Besides exceptions (Kappel & Das, 2019; Delgado Blanco et al., 2019), traditional approaches for the prediction of RNA-protein complexes can generally be roughly divided into two groups, free docking approaches (Pérez-Cano et al., 2010; Setny & Zacharias, 2011; Huang et al., 2013; Guilhot-Gaudeffroy et al., 2014; Tuszynska et al., 2015; Iwakiri et al., 2016; Van Zundert et al., 2016; Arnautova et al., 2018) and template-based docking approaches (Zheng et al., 2016; Zhang et al., 2022). Here, we focus on deep learning based methods for RNA-protein complex predictions, but refer the interested reader to some excellent reviews of the field (Nithin et al., 2018; Bheemireddy et al., 2022; Liu et al., 2023).

### Deep Learning based RNA-Protein Structure Prediction

After the remarkable success of AlphaFold 2 (Jumper et al., 2021), predicting protein structures with nearly experimental accuracy, and its extension AlphaFold Multimer (Evans et al., 2021) for the prediction of protein multimers, recent deep learning approaches for the prediction of the structure of biological macromolecules extended their capabilities to other molecular entities like DNA, RNA, and small molecule ligands (Abramson et al., 2024; Krishna et al., 2024; Baek et al., 2024; Roche et al., 2024). However, while the quality of protein predictions is typically retained, particularly modeling RNAs remains challenging (Das et al., 2023; Bernard et al., 2025). This also transfers to predictions of RNA-protein interactions, and new algorithms that can improve the interaction prediction quality are highly sought after.

### Diffusion Models

Diffusion generative models (DGMs) have emerged as a powerful framework for modeling complex probability distributions, offering advantages over traditional likelihood-based and implicit generative approaches (Ketata et al., 2023). DGMs operate by defining a diffusion process that gradually transforms the data distribution into a tractable prior. The key insight lies in learning the score function – the gradient of the log probability density function ∇**_x_** log *p_t_*(**x**)^2^ – of this evolving distribution. Once learned, this score function enables sampling from the underlying probability distribution through established algorithms (Song et al., 2020). The success of DGMs has led to their widespread adoption in computational biology. These applications span diverse tasks including conformer generation (Jing et al., 2022; Xu et al., 2022; YanWang et al., 2024; Fan et al., 2024; Park & Shen), molecule generation (Hoogeboom et al., 2022), RNA secondary structure generation (Wang et al., 2025), and protein design (Trippe et al., 2022; Liu et al., 2024). Particularly noteworthy are their contributions to protein structure (Jing et al.; Watson et al., 2023; Wu et al., 2024) and backbone generation (Yim et al., 2023) as well as protein-protein docking (Ketata et al., 2023), where they have demonstrated remarkable capabilities in capturing complex structural relationships. AlphaFold 3 utilizes a diffusion-based architecture to predict the structures of various biomolecular complexes, including proteins, nucleic acids, and small molecule ligands (Abramson et al., 2024).

In this work, we employ a new strategy that leverages the recent advancements of deep learning models for structural biology by directly utilizing their predictions. However, in contrast to methods like ProRNA3D-single (Roche et al., 2024), which is trained using RNA monomer predictions from RhoFold (Shen et al., 2024) and protein monomer predictions from ESMFold (Lin et al., 2023), STRAND is trained directly on RNA-protein complexes of experimentally validated structures from the PDB. This allows us to use any RNA-protein structure as a starting point for the backward diffusion process, including predictions, making it independent of the underlying structure generation method at test time.

## 3. Methods

In this section, we describe STRAND, our extension of DiffDock-PP (Ketata et al., 2023) to model monomeric RNA-protein complexes. Similar to Ketata et al. (2023), we model proteins and RNA at the residue level in STRAND. For the protein, we exactly follow Ketata et al. (2023), using the same node features as used in DiffDock-PP that represent each residue by its type, the position of its *α*-carbon atom, and the processed embeddings from ESM2 (Lin et al., 2023). To enable processing of RNAs, we use similar node features: the nucleotide type, the coordinates of the phosphate atoms to represent the backbone, and the embeddings obtained from running RNA-FM (Chen et al., 2022). For the diffusion process, we treat one molecular entity as the receptor (the RNA) and apply noise to the other one (the protein). In the following, we describe STRAND in more detail with a focus on differences compared to DiffDock-PP.

### 3.1. STRAND

We denote **X**_1_ ∈ ℝ^3*n*^ as the ligand consisting of *n* residues and **X**_2_ ∈ ℝ^3*m*^ as the receptor with *m* residues. Ketata et al. (2023) assigns the ligand/receptor based on residue length. In contrast, we run the diffusion process on the protein, regardless of the relative sizes. While this increases training time due to longer molecules being diffused, it improves stability during evaluation, where test set lengths differ from those in training, thus we assign the protein **X**_1_ as the lig-and. With 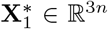 and 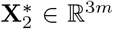 denoting the initial positions in space of both ligand and receptor respectively, the receptor is kept fixed 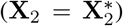, and the task is to predict the position of the ligand with respect to the receptor. We define RNA-protein structure refinement as learning the conditional probability distribution *p*(**X**_1_ | **X**_2_), which describes the possible ligand poses within a submanifold ℳ, given the receptor structure **X_2_**. Following Ketata et al. (2023), we avoid the inefficiencies arising from learning DGMs on arbitrary submanifolds (Bortoli et al., 2022), using the framework of intrinsic diffusion models (Corso et al., 2022).

#### Backbone Refinement via Diffusion

Building on the framework of Ketata et al. (2023), which is limited to rigidbody transformations, we extend the approach by training separate models for translation, rotation, torsion, and the combinations rotation and translation, and rotation, translation, and torsion.

Formally, we introduce the 3D translation group *T* (3), the 3D rotation group *SO*(3), and associate changes in torsion angles at each rotatable bond with a copy of the 2D rotation group *SO*(2).

The translation operation *A*_tr_ : *T* (3) *×* ℝ^3*n*^→ ℝ^3*n*^ is naturally defined as:

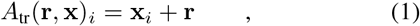

where **x***_i_* ∈ ℝ^3^ denotes the position of the *i*th backbone residue.

The rotation operation *A*_rot_ : *SO*(3) *×* ℝ^3*n*^ → ℝ^3*n*^ is defined as:

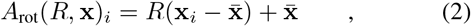

where 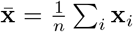 is the (unweighted) center of mass of the ligand. This corresponds to a rotation around the center of mass.

For torsion, we follow the definition introduced in Diff-Dock (Corso et al., 2022), where changes in torsion angles are disentangled from global rotations and translations. In DiffDock’s approach, the ambiguity of torsion changes, where the torsion angle around any bond (*a_i_, b_i_*) could be updated by rotating the *a_i_* side, the *b_i_* side, or both, is resolved by defining the action of elements of *SO*(2)*^m^* to cause minimal perturbation (in an RMSD sense) to the structure.

More precisely, let *B_k_, θ_k_*(**x**) ∈ ℝ^3*n*^ be any valid torsion update by *θ_k_* around the *k*th rotatable bond (*a_k_, b_k_*). We define *A*_tor_ : *SO*(2)*^m^ ×* ℝ^3*n*^ → ℝ^3*n*^ such that

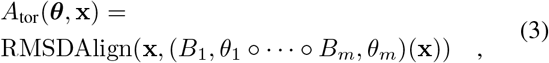

where ***θ*** = (*θ*_1_, …, *θ_m_*), and

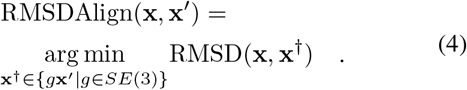

In words, all *m* torsion updates are applied in any order before global RMSD alignment with the unmodified pose.

Now consider the product space ℙ = *T* (3) *× SO*(3) *× SO*(2)*^m^*, and define *A* : ℙ *×* ℝ^3*n*^ → ℝ^3*n*^ as:

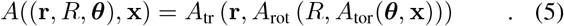

Using this formulation, the submanifold of ligand poses can be described as:

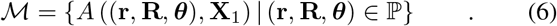

Following Corso et al. (2022), each mapping *A*(·, **X**_1_) is a bijection, ensuring the existence of inverse maps. This allows us to define a diffusion process over the full product space P to model the ligand pose distribution. In scenarios where the model is trained specifically for rotation, we set the translation vector **r** ∈ *T* (3) to the zero vector. Conversely, when training for translation only, we set the rotation matrix **R** ∈ *SO*(3) to the identity matrix **I**, indicating the absence of rotation. When torsion is not modeled, we restrict the ligand to be rigid by eliminating any rotatable bonds.

Since each component of ℙ forms a product manifold, we define independent forward diffusion processes (Rodola et al., 2019), where the score lies in the corresponding tangent space (Bortoli et al., 2022). The forward SDE is given by:

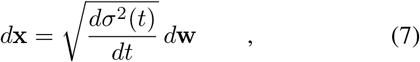

where 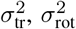, and 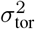 govern diffusion in *T* (3), *SO*(3), and *SO*(2)*^m^*, respectively. Their joint effect governs diffusion in ℙ. The term *d***w** denotes Brownian motion on the respective manifolds.

#### Neural Architecture

We adopt the architecture proposed by Ketata et al. (2023), originally developed for protein–protein complexes. To account for the differences in molecular composition and interaction dynamics between protein–protein and protein–RNA complexes, we introduce separate embedding layers for protein and RNA components. Furthermore, we extend the original model to additionally predict torsional scores, enabling finer-grained structural modeling.

#### Training Details

For training, we use all RNA and protein containing PDB samples with deposition date before September 30, 2021, the training cutoff date of AlphaFold 3, downloaded on January 3, 2025. We then use a preprocessing pipeline to obtain only interacting monomeric RNA and protein chains. While the natural choice would be to collect monomeric RNA-protein complexes only, we found that adding RNA-protein multimers during training and using all chains of one molecular type as the receptor and all others together as the ligand is beneficial during training for all STRAND versions except STRAND_tr+rot+tor_, our version using all transformations. We speculate that this process might help to directly learn the interactions of all chains in a single process. We train five different models for backbone refinement, one for translation only (STRAND_tr_), one for rotation only (STRAND_rot_), one for torsion only (STRAND_tor_), and the combinations of translation and rotation (STRAND_tr+rot_), as well as translation, rotation and torsion (STRAND_tr+rot+tor_). The models are trained on a single A40 GPU for roughly 5.75, 33, 6, 72, and 54 hours, respectively, using Adam. Please find details about hyperparameter settings in Appendix A.

## 4. Experiments

To assess STRAND’s refinement capabilities, we initialize STRAND with the predictions of two recently proposed approaches for the modeling of monomeric RNA-protein complexes, AlphaFold 3 (Abramson et al., 2024) and ProRNA3D-single (Roche et al., 2024), and sample 40 new poses from the different models. For AlphaFold 3, we always use sample-0 predictions as only small variance is reported for the sampling from the diffusion model in Al-phaFold 3 (Abramson et al., 2024), which is in line with our preliminary experiments, where we also did not observe strong variance across seeds nor different samples of AlphaFold 3. Predicted sequences are aligned to the ground truth via Needleman-Wunsch (Needleman & Wunsch, 1970) to handle misalignments. We report performance in terms of complex RMSD (cRMSD), determined by superimposing the ground truth and predicted complex structures via the Kabsch algorithm (Kabsch, 1976) and computing the RMSD between all C*α* and P coordinates following Ganea et al. (2021).

### Data

For all our experiments with AlphaFold 3, we use a randomly selected set of monomeric RNA-protein complexes published in PDB after the AlphaFold 3 training cutoff date to ensure independence from the training set. We split the samples based on experimental techniques into a set of test structures derived from X-ray crystallography and one with samples derived by other techniques like NMR and cryo EM, and evaluate the refinement of AlphaFold 3 predictions on both splits. The datasets contain 35 and 46 samples for X-ray and non-X-ray, respectively. For the refinement of ProRNA3D-single predictions, we only use data from X-ray crystallography, but download an additional set of 28 complexes from the PDB. The X-ray dataset for evaluations with ProRNA3D-single then contains a total of 63 samples. For more details about the datasets, please see Appendix B.

### 4.1. RNA-protein Complex Refinement with STRAND

In this section, we evaluate different STRAND models trained for different transformations to assess their influence on performance: the translation only model STRAND_tr_, the rotation only model STRAND_rot_, the torsion only model STRAND_tor_, and the combined versions STRAND_tr+rot_ and STRAND_tr+rot+tor_. As described in Section 3, we diffuse over the protein, while the RNA is treated as the receptor. We sample 40 poses for each task for each individual model. The best sample is selected manually based on the complex RMSD (cRMSD).

#### 4.1.1. STRAND canImproveAlphaFold3

We summarize the results for the AlphaFold 3 predictions for the X-ray and non-X-ray derived monomeric complexes from PDB and their refinement with STRAND in Table 1.

**Table 1.** RNA-protein backbone refinement results of STRAND for AlphaFold 3 predictions. We show results for the translation only model STRAND_tr_, the rotation only model STRAND_rot_, the torsion only model STRAND_tor_, and the combined versions STRAND_tr+rot_ and STRAND_tr+rot+tor_. Best performance is indicated in bold, equal best performance is additionally indicated by underlined scores. Results are reported in terms of mean and median complex RMSD (cRMSD; lower is better) and percentage of samples below a certain cRMSD threshold (2Å, 5Å, 10Å ; higher is better). We observe that STRAND_tr+rot_ shows the best performance on average, while at least one of the STRAND models performs equally well or better than AlphaFold 3 across all metrics for both datasets. The worst performance is observed for the models trained with torsion.

| Methods | Complex RMSD (Å) ↓ |  |  |  |  |  |  |  |  |  |
| --- | --- | --- | --- | --- | --- | --- | --- | --- | --- | --- |
|  | X-Ray |  |  |  |  | Non X-Ray |  |  |  |  |
|  | %<2Å | %<5Å | %<10Å | Median | Mean | %<2Å | %<5Å | %<10Å | Median | Mean |
| AlphaFold 3 | 34.29 | 85.71 | **100.00** | 2.20 | 3.09 | 19.57 | 45.65 | **63.04** | 7.15 | 10.86 |
| STRAND<sub>tr</sub> | 34.29 | 82.86 | **100.00** | 2.30 | 3.06 | 19.57 | **47.83** | 60.87 | **6.53** | 10.71 |
| STRAND<sub>rot</sub> | 37.14 | **88.57** | 97.14 | 2.23 | 3.07 | **21.74** | 43.48 | **63.04** | 6.80 | 10.69 |
| STRAND<sub>tor</sub> | 28.57 | 85.71 | **100.00** | 2.71 | 3.29 | 4.35 | 41.30 | **63.04** | 7.13 | 11.08 |
| STRAND<sub>tr+rot</sub> | **42.86** | 82.86 | **100.00** | **2.16** | **2.97** | 17.39 | 45.65 | 60.87 | 6.87 | **10.41** |
| STRAND<sub>tr+rot+tor</sub> | 31.43 | 82.86 | **100.00** | 3.10 | 3.45 | 0.00 | 34.78 | 54.35 | 8.73 | 11.64 |

##### Single Transformations

While AlphaFold 3 shows remarkable prediction quality on the X-ray data with a mean cRMSD of 3.09 and 100% of the predictions being below a cRMSD of 10Å, we still see some improvements when applying STRAND to these strong predictions. For the models trained for single transformations, two out of three models improve the mean performance (STRAND_tr_ with 3.06Å and STRAND_rot_ with 3.07Å) while the model trained with torsion achieves a slightly higher mean cRMSD of 3.29Å. In addition, we observe remarkable increases by 2.85% of the percentage of complexes predicted below a cRMSD of 2Å and by 2.86% for complexes below 5Å, with only slightly worse median performance (2.23Å and 2.20Å for STRAND_rot_ and AlphaFold 3, respectively) for structure refinement done via STRAND_rot_. Generally, we observe improvements for 48.5% and 40% of the X-ray modeling tasks for the individual models STRAND_tr_ and STRAND_rot_, respectively (see also Figures 2, 3, and 4 for results on individual tasks).

**Figure 2.**
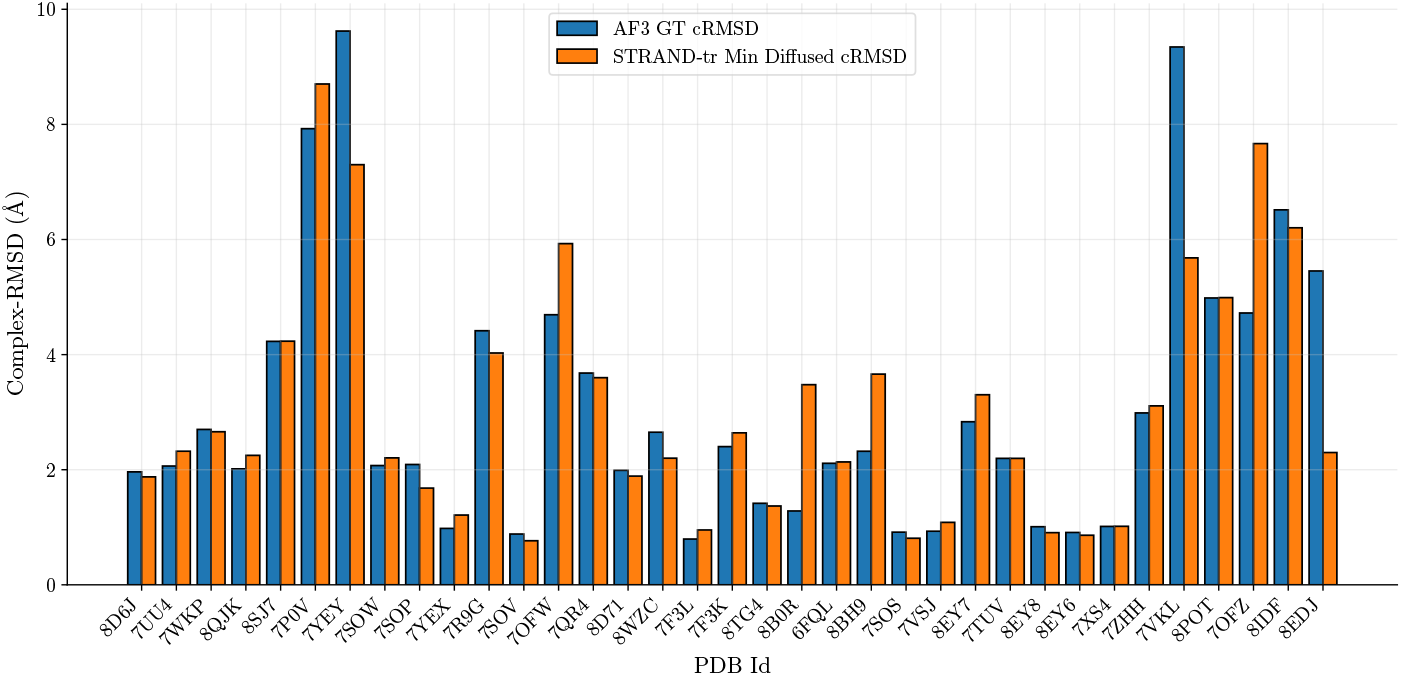
Per sample cRMSD of AlphaFold 3 predictions on X-ray data and the corresponding prediction of STRAND. We show results of the translation-only model STRAND_tr_.

**Figure 3.**
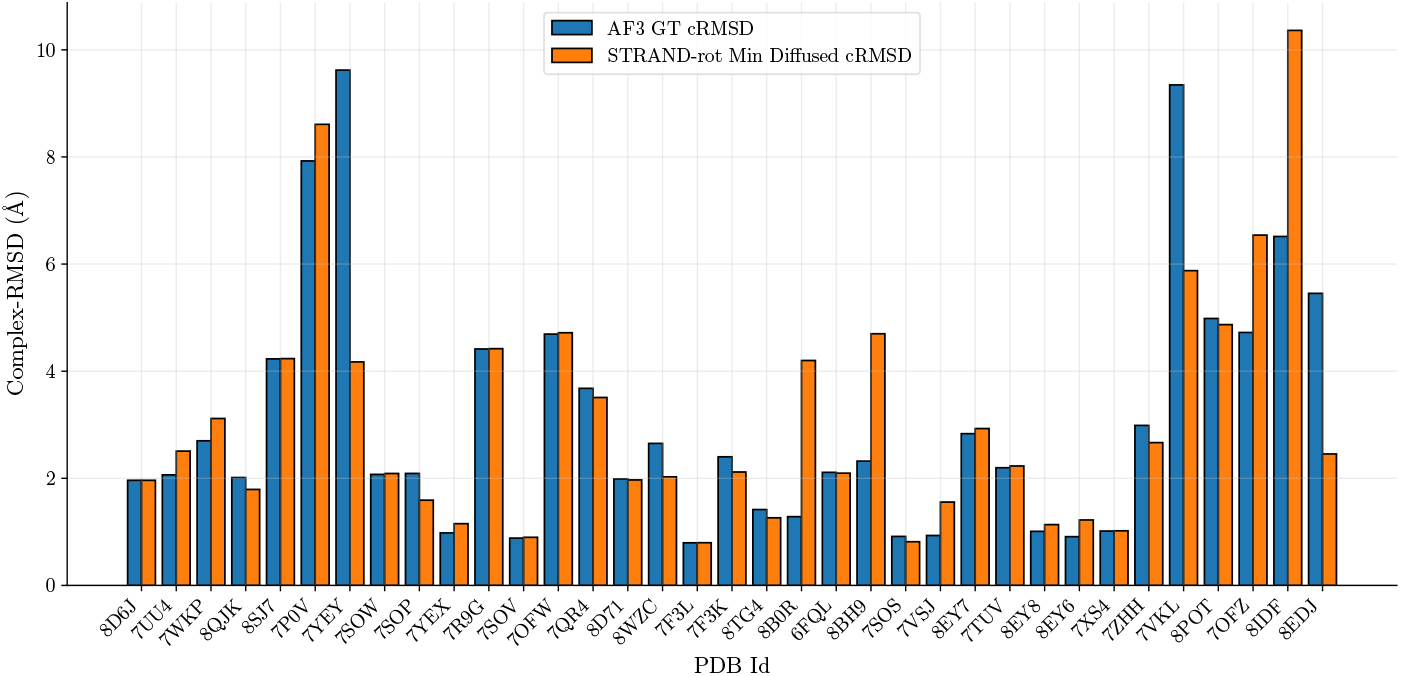
Per sample cRMSD of AlphaFold 3 predictions on X-ray data and the corresponding prediction of STRAND. We show results of the rotation-only model STRAND_rot_.

**Figure 4.**
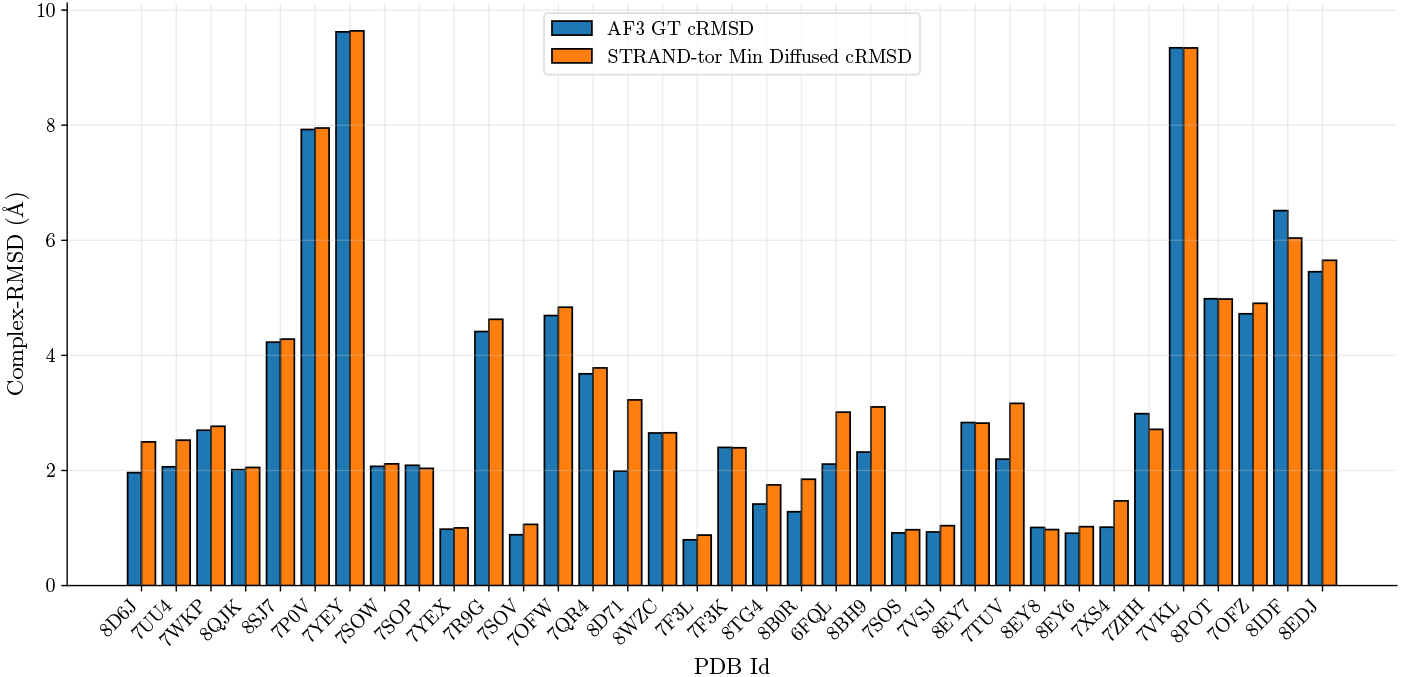
Per sample cRMSD of AlphaFold 3 predictions on X-ray data and the corresponding prediction of STRAND. We show results of the torsion-only model STRAND_tor_.

**Figure 5.**
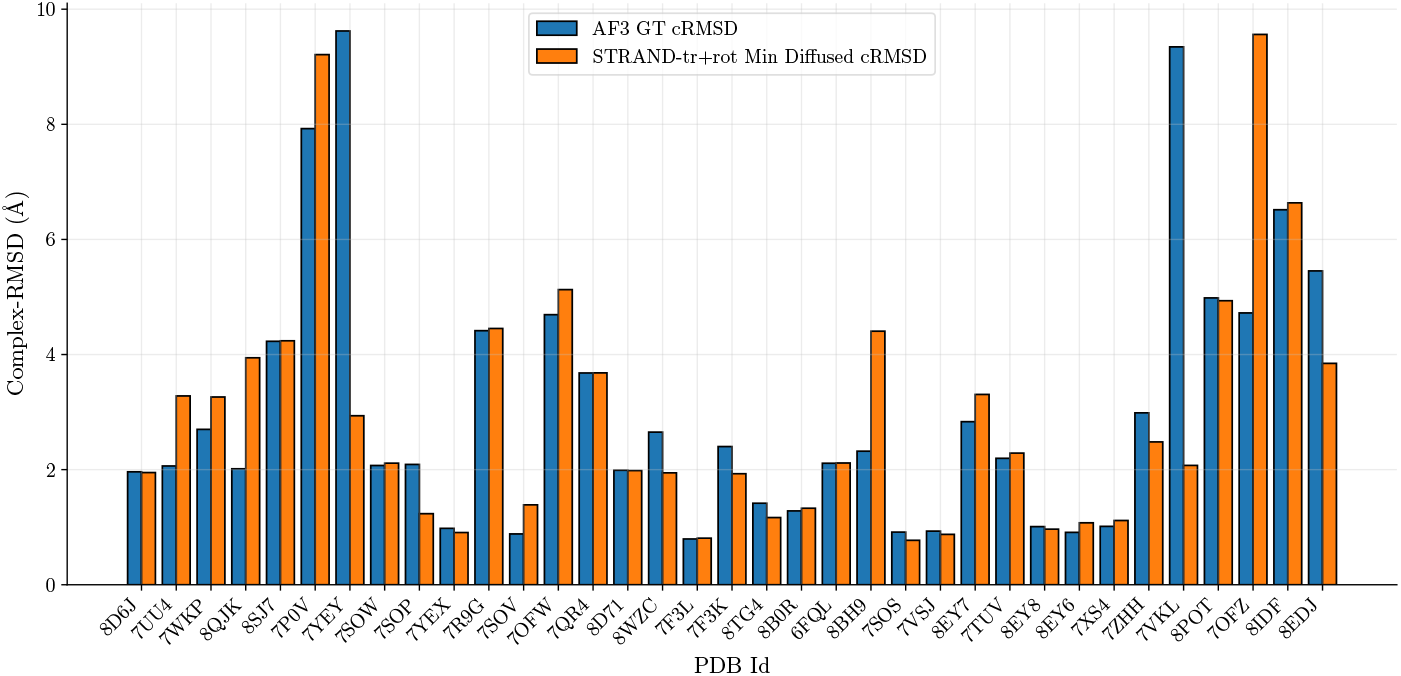
Per sample cRMSD of AlphaFold 3 predictions on X-ray data and the corresponding prediction of STRAND. We show results of the combined model of translation and rotation STRAND_tr+rot_.

**Figure 6.**
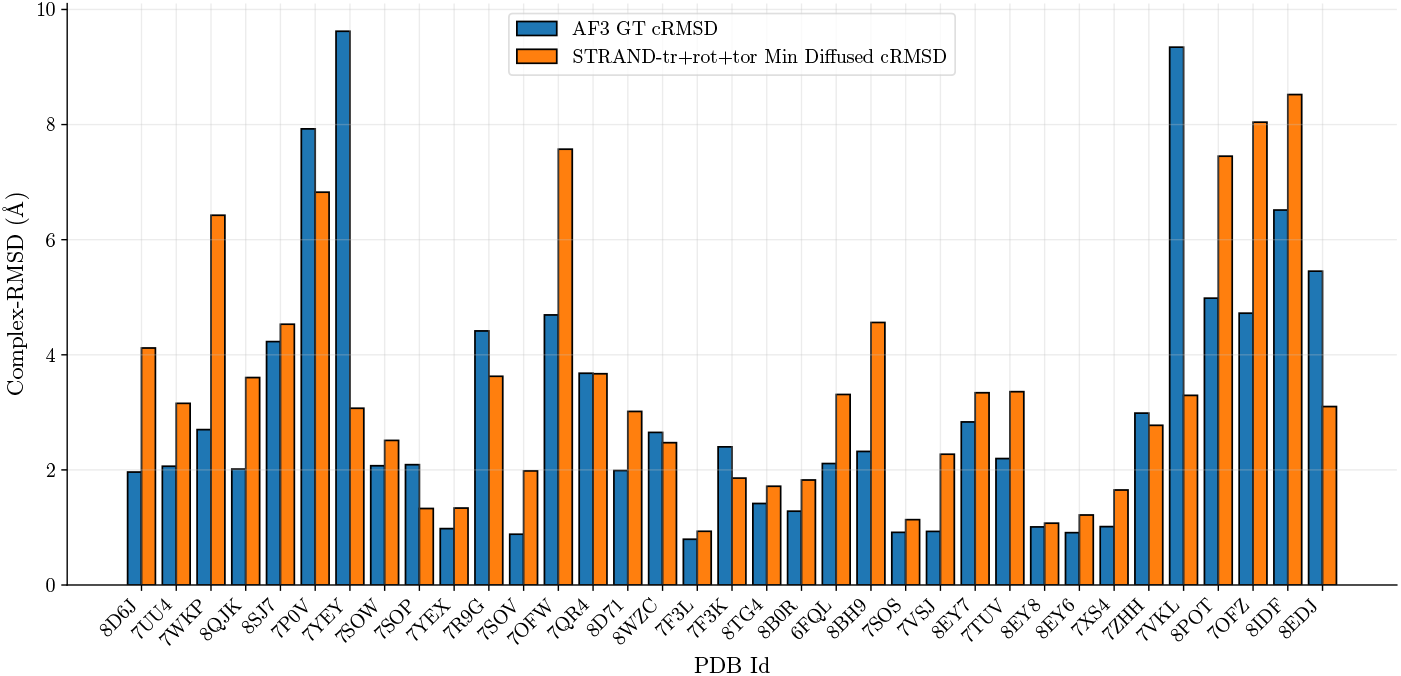
Per sample cRMSD of AlphaFold 3 predictions on X-ray data and the corresponding prediction of STRAND. We show results of the combined model for translation, rotation, and torsion STRAND_tr+rot+tor_.

For the non-X-ray data, we generally observe worse predictions of AlphaFold 3 with only roughly 60% of the predictions being below a cRMSD of 10Å. However, the trends observed for the different STRAND versions are also confirmed for non-X-ray samples. Again, STRAND_rot_ achieves the highest percentage of predictions below 2Å (2.17% improvement) and achieves the lowest mean cRMSD of all single transformation models (10.69Å for STRAND*_rot_* compared to 10.86Å for AlphaFold 3) while the torsion only model shows the highest mean cRMSD of 11.08Å. Remarkably, the model trained with translation only achieves the highest percentage of predictions below a cRMSD of 5Å, a 2.18% improvement over AlphaFold 3. Overall, the three models can improve 63%, 63% and 26% of the individual tasks for STRAND_tr_, STRAND_rot_, and STRAND_tor_, respectively. We show visualizations for the performance on individual samples in Figures 7, 8, and 9 in Appendix C.2.

**Figure 7.**
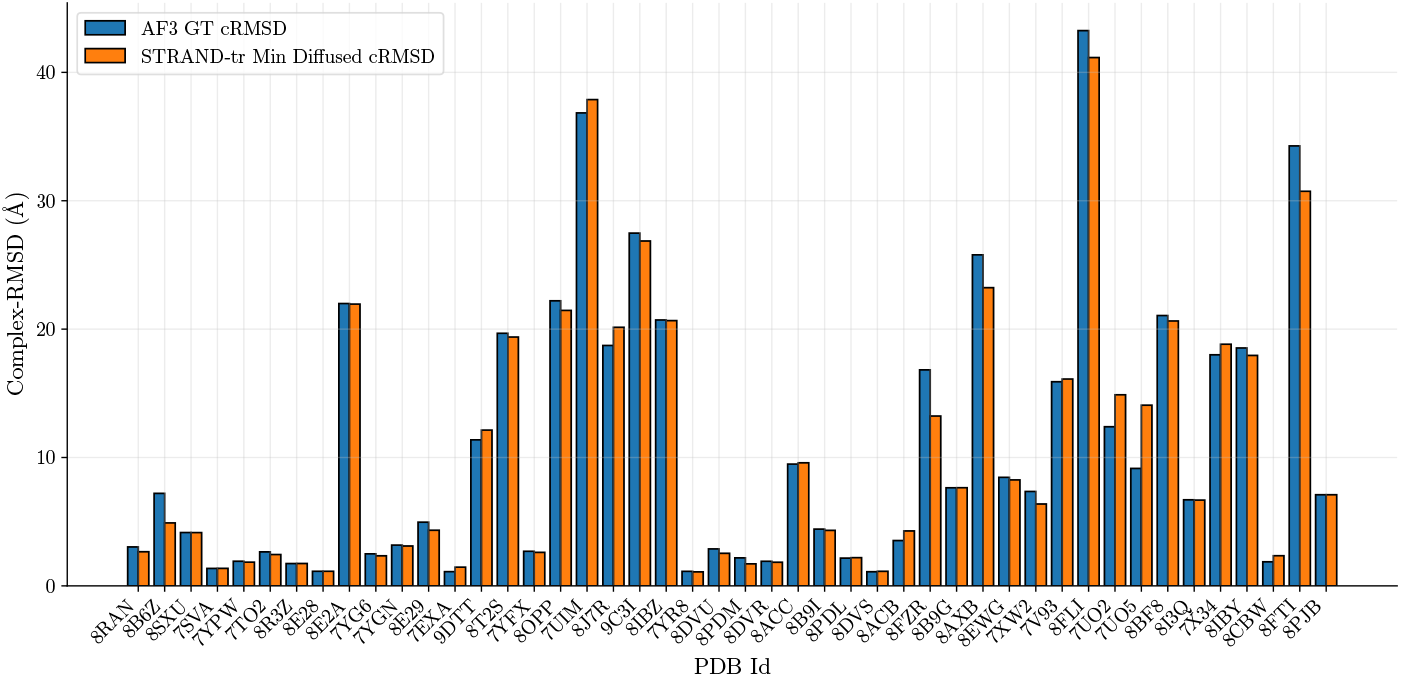
Per sample cRMSD of AlphaFold 3 predictions on non-X-ray data and the corresponding prediction of STRAND. We show results of the translation-only model STRAND_tr_.

**Figure 8.**
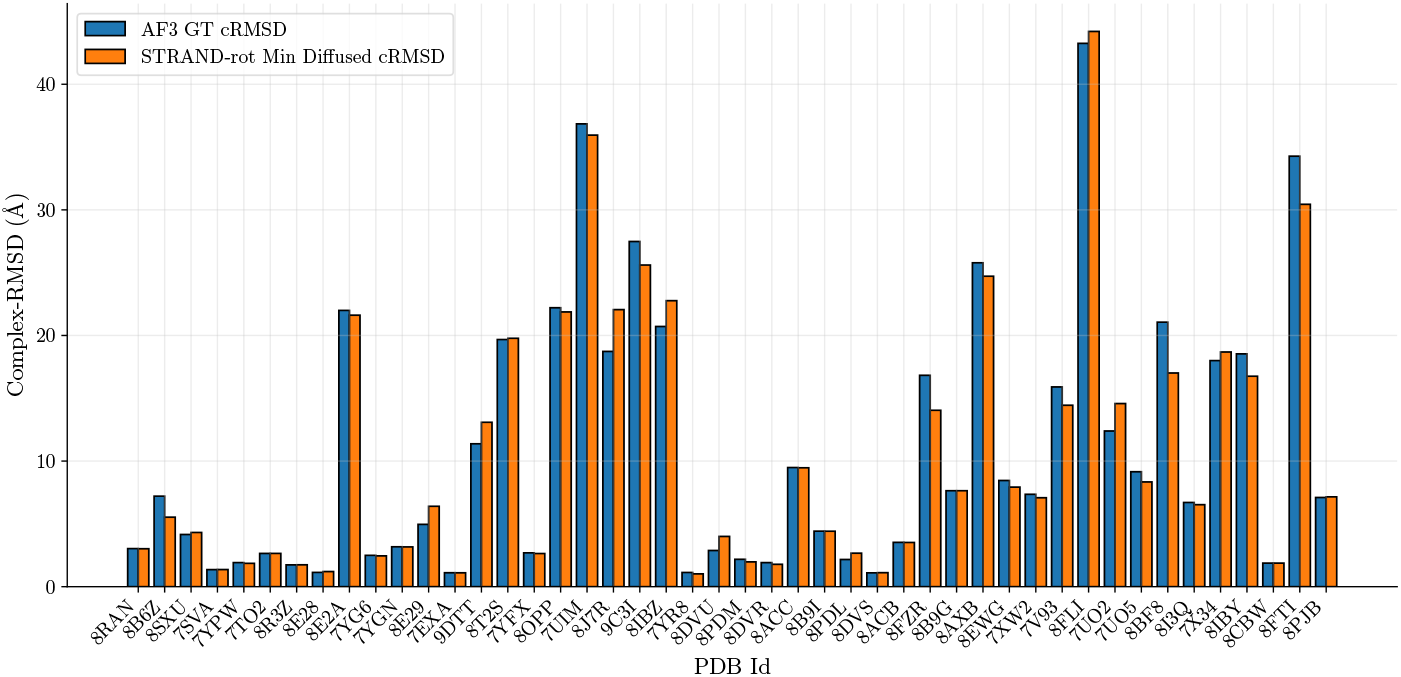
Per sample cRMSD of AlphaFold 3 predictions on non-X-ray data and the corresponding prediction of STRAND. We show results of the rotation-only model STRAND_rot_.

**Figure 9.**
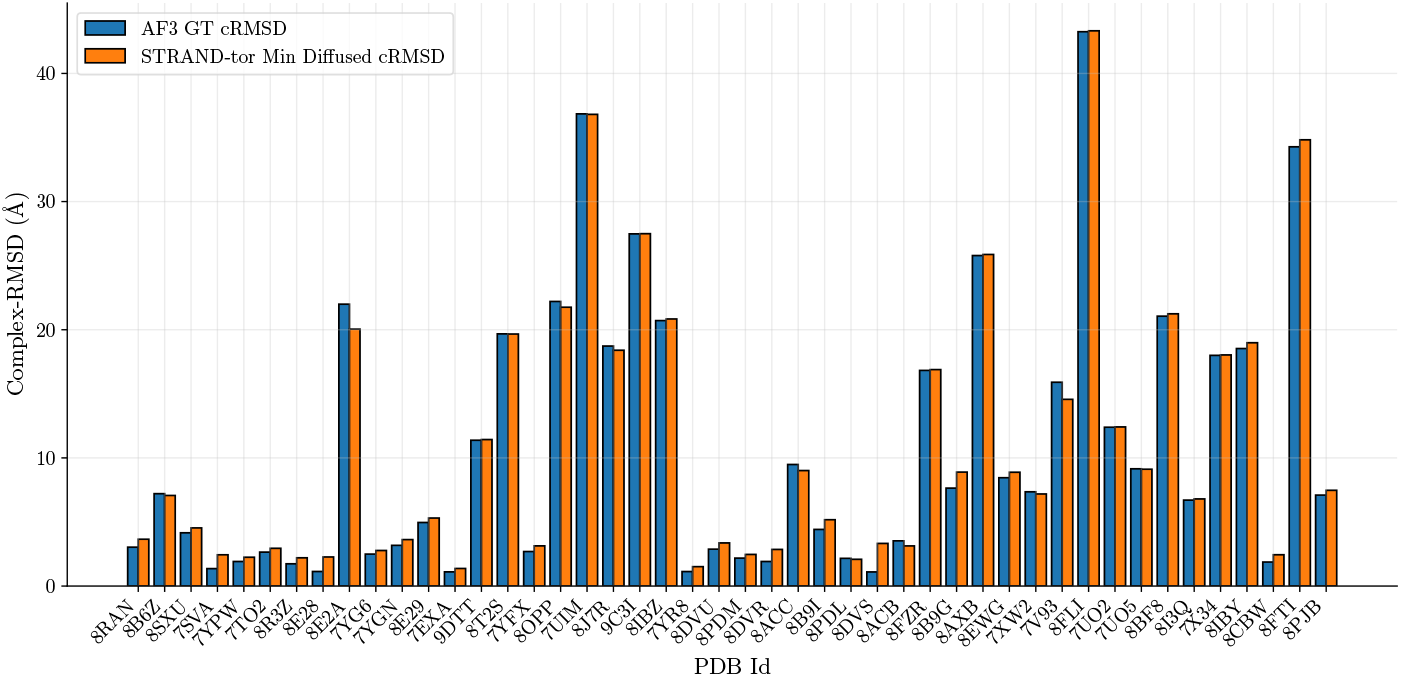
Per sample cRMSD of AlphaFold 3 predictions on non-X-ray data and the corresponding prediction of STRAND. We show results of the torsion only model STRAND_tor_.

**Figure 10.**
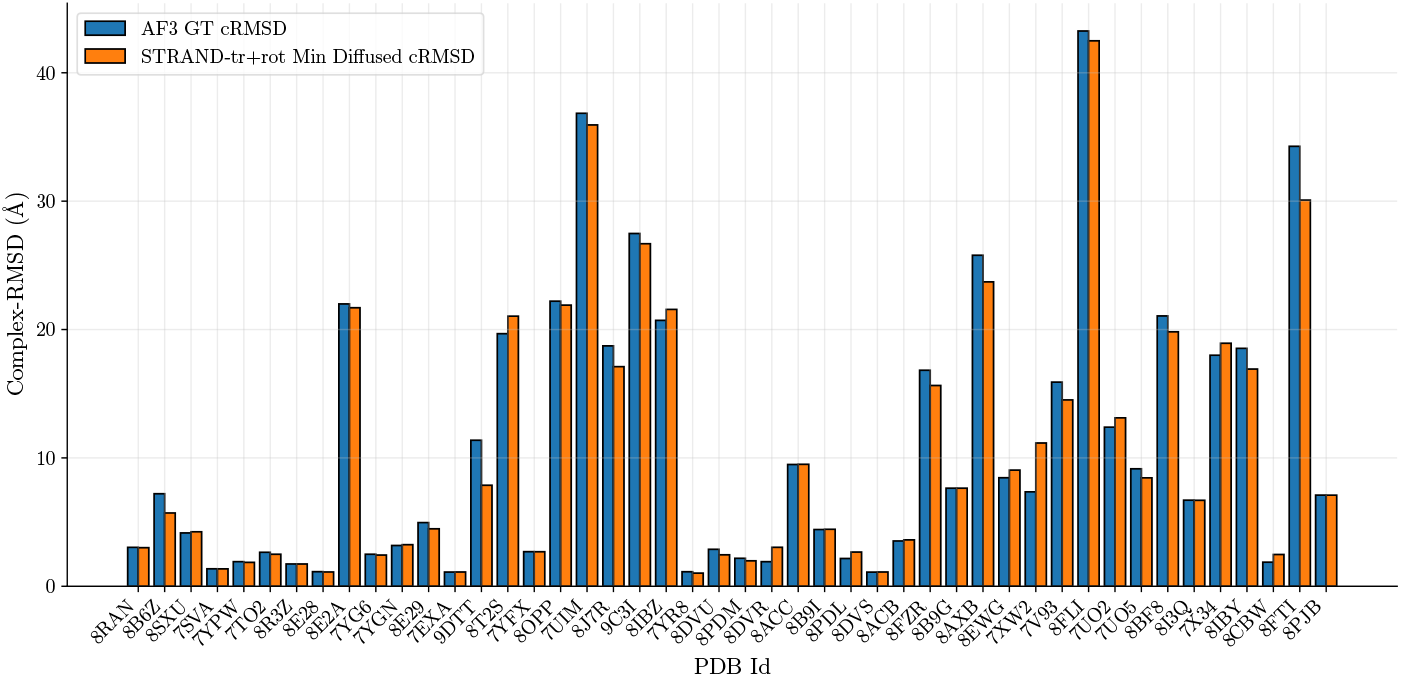
Per sample cRMSD of AlphaFold 3 predictions on non-X-ray data and the corresponding prediction of STRAND. We show results of the combined model of translation and rotation STRAND_tr+rot_.

**Figure 11.**
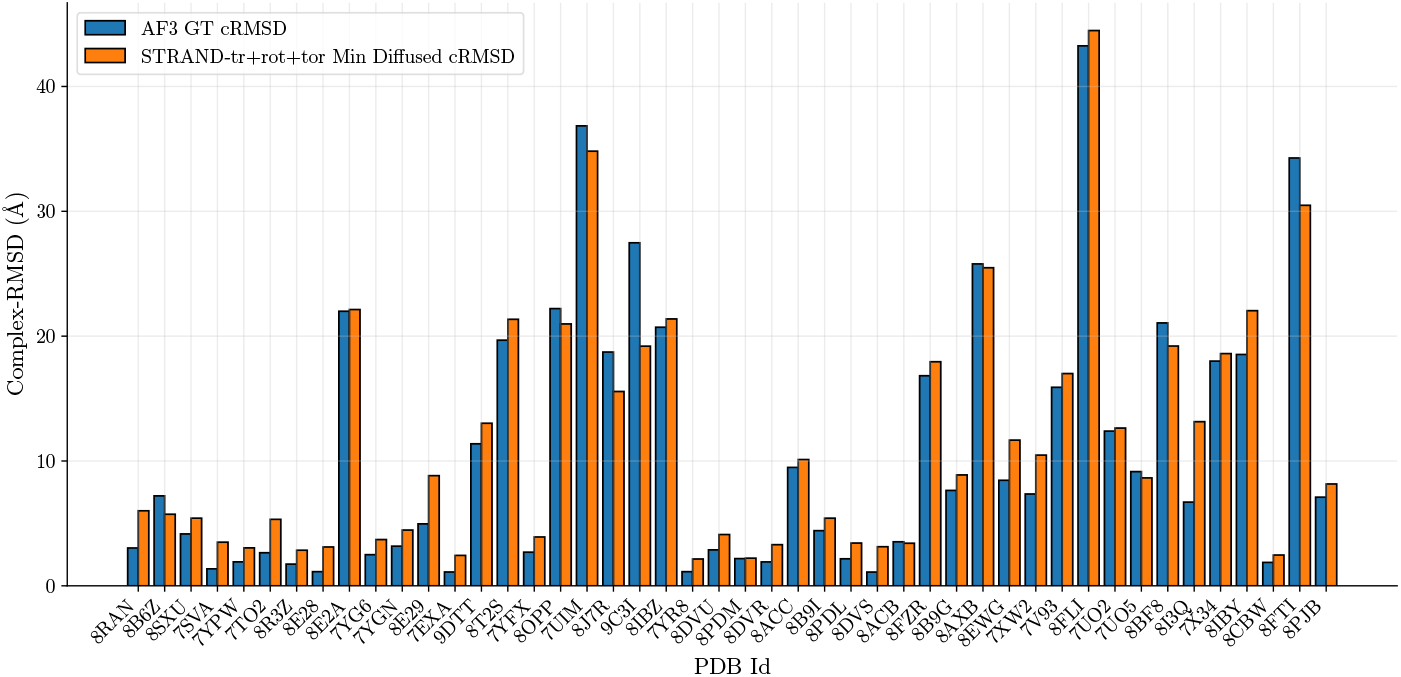
Per sample cRMSD of AlphaFold 3 predictions on non-X-ray data and the corresponding prediction of STRAND. We show results of the combined model of translation, rotation, and torsion STRAND_tr+rot+tor_.

**Figure 12.**
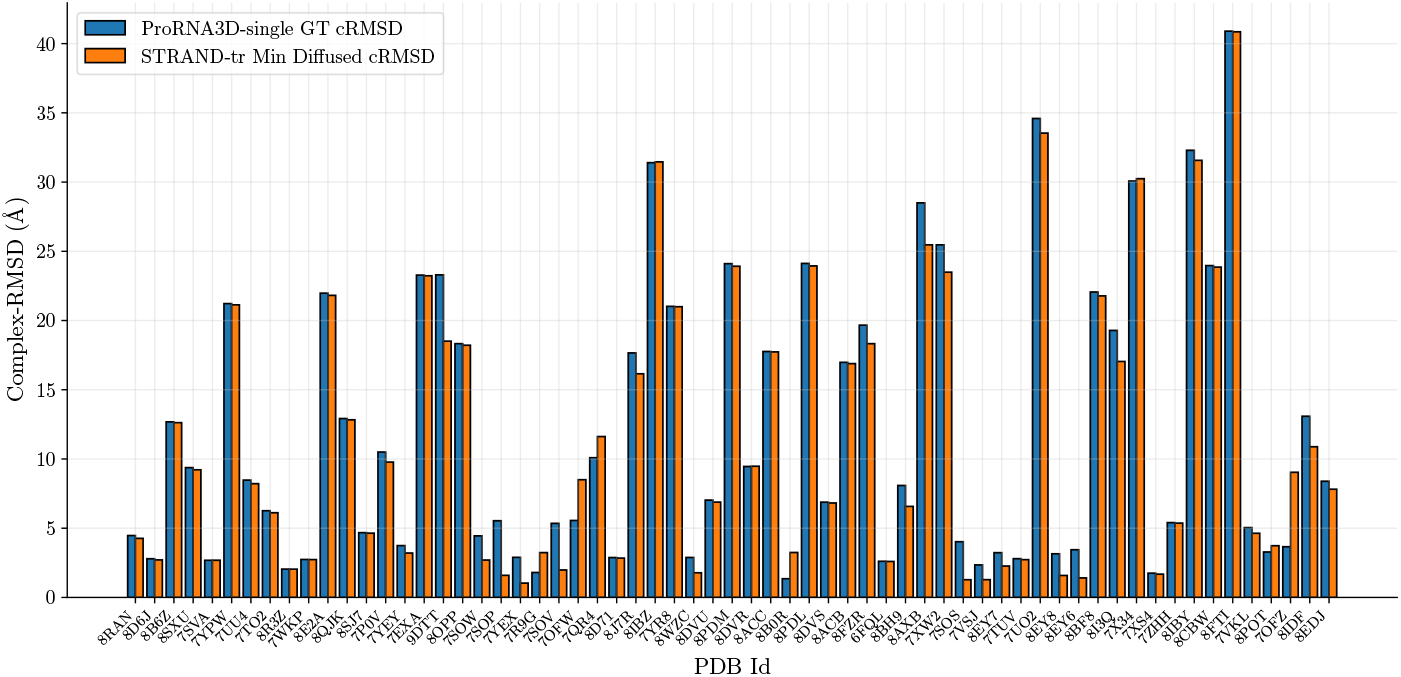
Per sample cRMSD of ProRNA3D-single predictions on X-ray data and the corresponding prediction of STRAND. We show results of the translation-only model STRAND_tr_.

**Figure 13.**
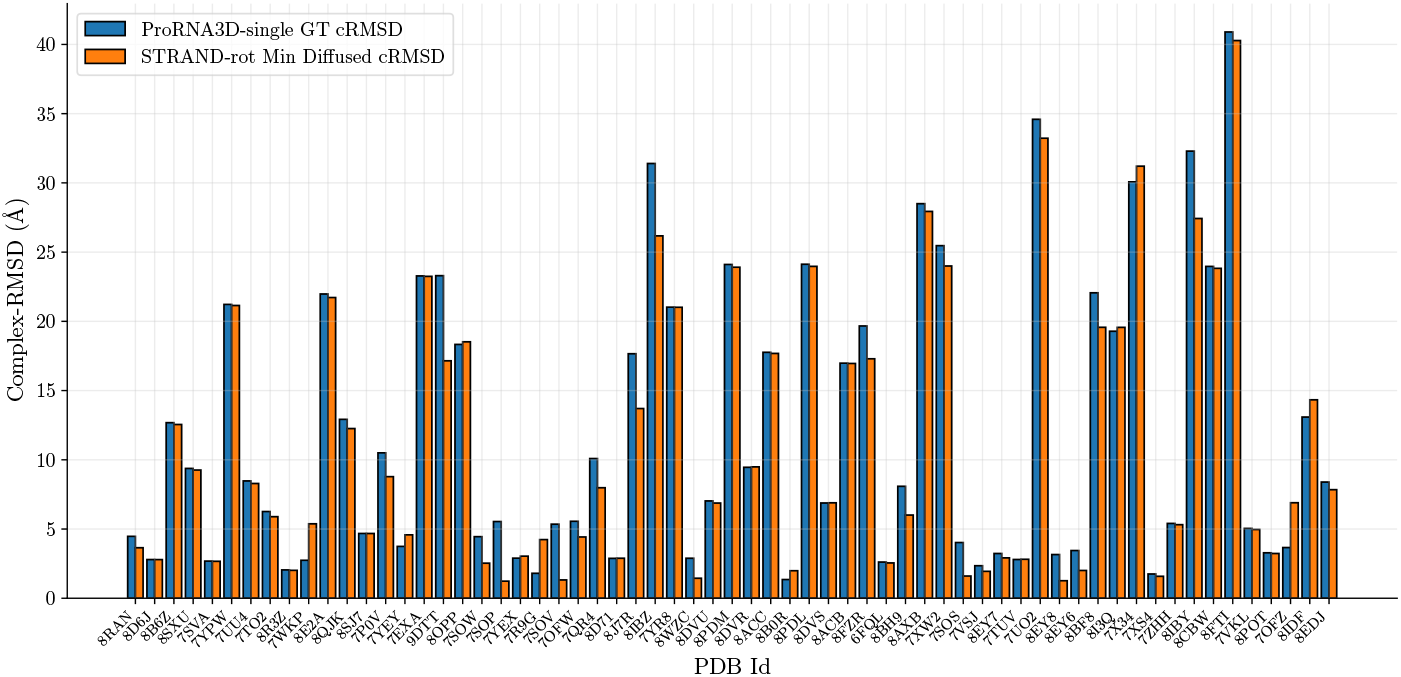
Per sample cRMSD of ProRNA3D-single predictions on X-ray data and the corresponding prediction of STRAND. We show results of the rotation-only model STRAND_rot_.

**Figure 14.**
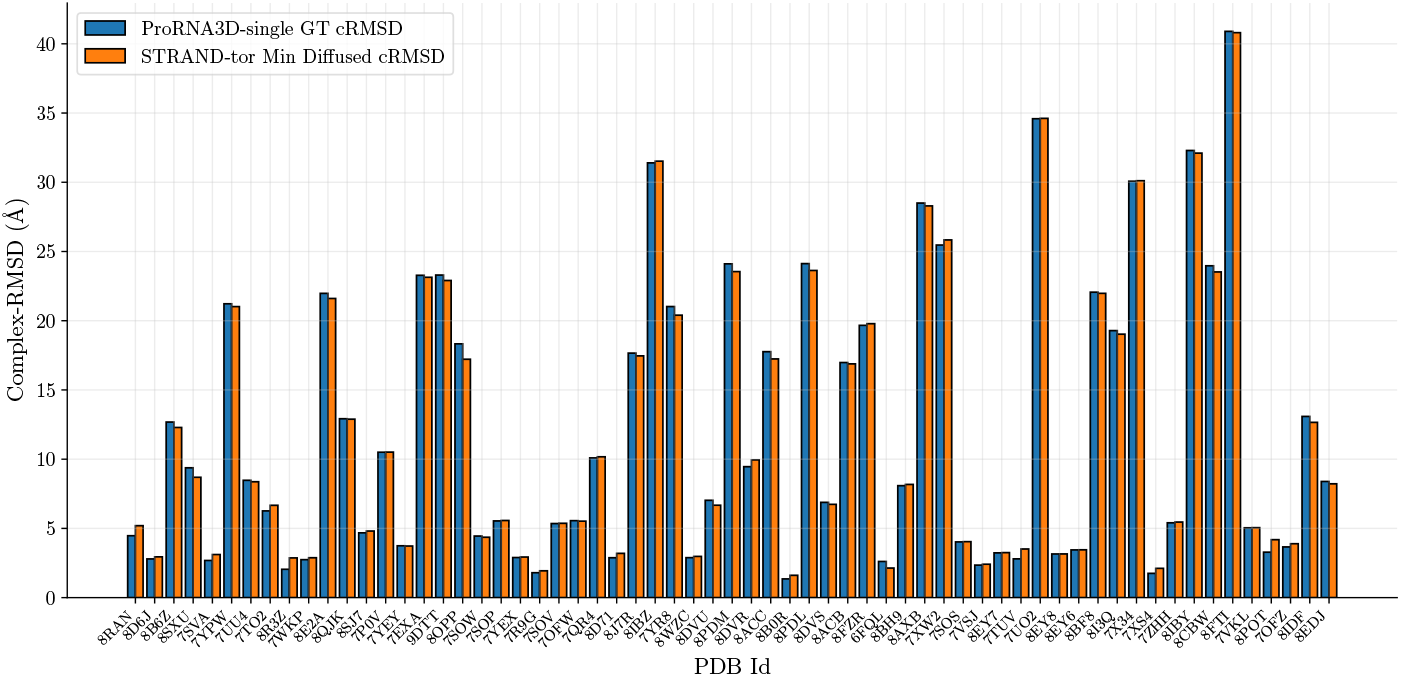
Per sample cRMSD of ProRNA3D-single predictions on X-ray data and the corresponding prediction of STRAND. We show results of the torsion-only model STRAND_tor_.

**Figure 15.**
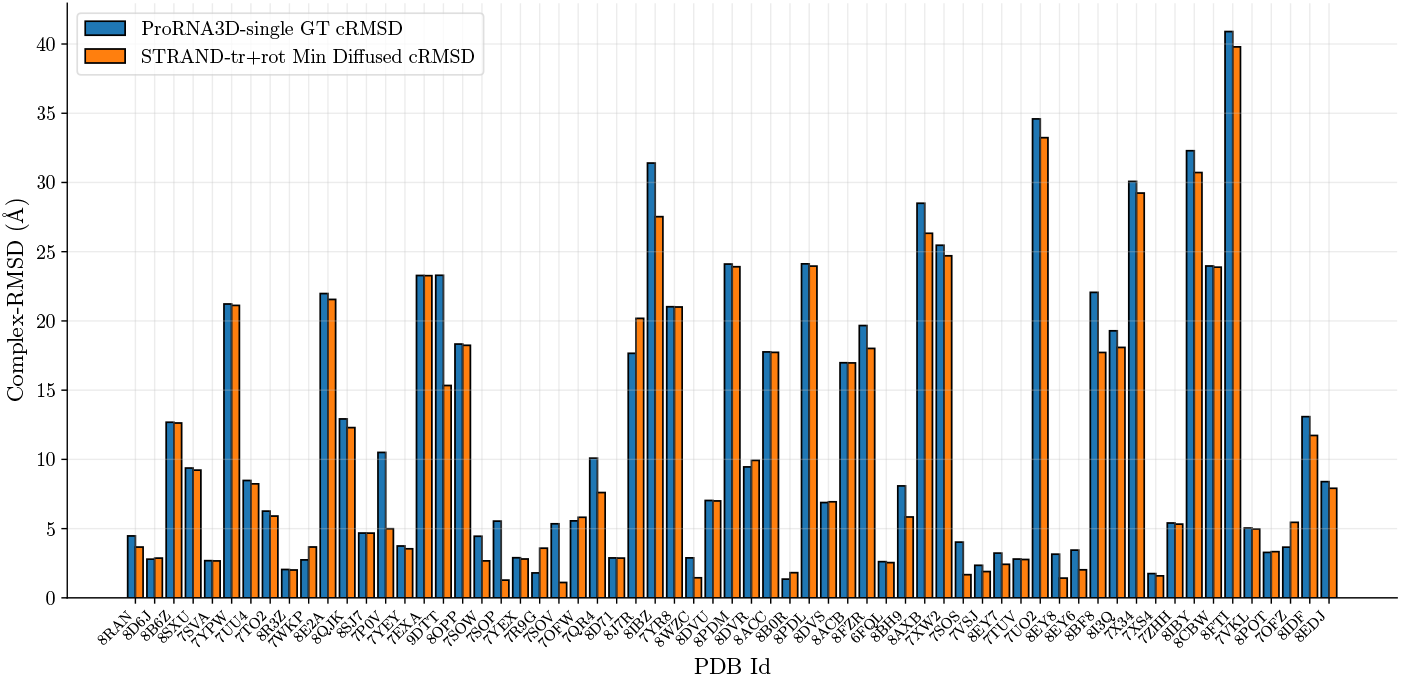
Per sample cRMSD of ProRNA3D-single predictions on X-ray data and the corresponding prediction of STRAND. We show results of the combined model for translation and rotation STRAND_tr+rot_.

**Figure 16.**
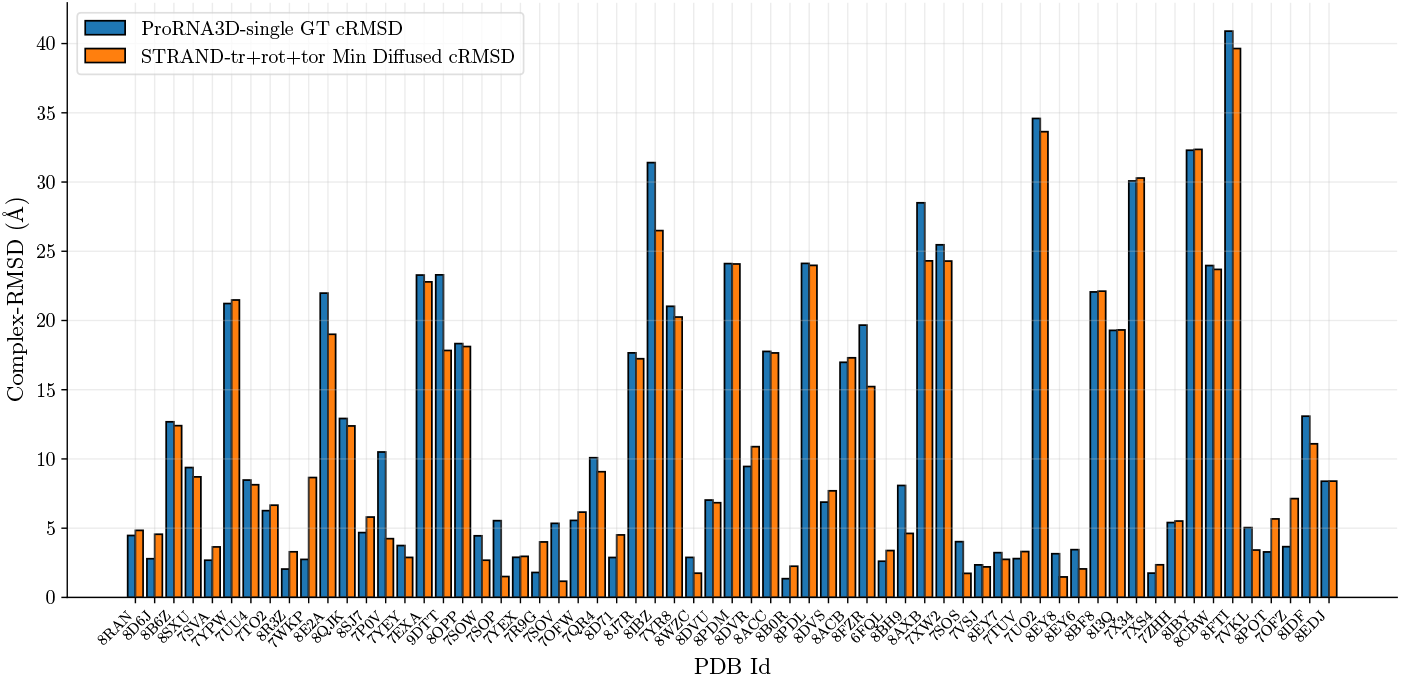
Per sample cRMSD of ProRNA3D-single predictions on X-ray data and the corresponding prediction of STRAND. We show results of the combined model for translation, rotation, and torsion STRAND_tr+rot+tor_.

Generally, the model trained only with torsion performs worse than the other two models. We speculate that the reason for this is that torsion only affects the internal RNA coordinates but not its general positioning with respect to the protein. Consequently, changes in overall complex RMSD through changes in the torsion angles can typically only be marginal. This can also be observed in Figure 4 and Figure 9 for the performance on individual samples, where the effect on cRMSD by changes in torsion angles is much smaller compared to the other transformations (compare e.g. with Figure 3) and further supported visually in Figure 1, where the large differences in cRMSD result from repositioning the two structures with respect to each other, while the local structures remain unchanged.

##### Combined Transformations

We also trained two models with combinations of transformations, STRAND_tr+rot_ trained with translation and rotation, and STRAND_tr+rot+tor_ trained with a combination of all the individual transformations. As shown in Table 1, we observe further improvements when combining the best-performing single transformations. Specifically, STRAND_tr+rot_ achieves the lowest mean cRMSDs of 2.97Å and 10.41Å for X-ray and non-X-ray samples, respectively. Furthermore, STRAND_tr+rot_ improves the percentage of samples below a cRMSD of 2Å by 8.57% and achieves also the lowest median cRMSD across all methods on the X-ray dataset. In contrast, the additional training for torsion seems to hurt performance in general, as STRAND_tr+rot+tor_ achieves the worst performance across all the different STRAND versions, while STRAND_tr+rot_ can improve 42% of the individual AlphaFold 3 predictions on the X-ray dataset and 63% of the AlphaFold predictions on the non-X-ray dataset. Please find plots for the performance on individual tasks for the combined models for the STRAND refinements of AlphaFold 3 predictions in Appendix C.1 and C.2.

#### 4.1.2. STRAND canImproveProRNA3D-single

To assess the performance of STRAND for a different predictor than AlphaFold 3, we decide to use the recently proposed ProRNA3D-single (Roche et al., 2024), which reports remarkable results for monomeric RNA-protein complex predictions. Please find an overview of the results for the X-ray dataset in Table 2. Remarkably, we observe that all versions of STRAND improve over ProRNA3D-single in terms of mean cRMSD. The general trend that training with translation, rotation, and the combination of both leads to the best results, as observed for STRAND refinement of AlphaFold 3 predictions, is also confirmed for ProRNA3Dsingle. However, the improvements appear more substantial: For predictions below a cRMSD of 2Å we observe an improvement of 9.53% and 7.94% for the translation only model and both the rotation and rotation+translation models, respectively. In addition, we find improvements for the predictions below 5Å and 10Å for all models except the torsion-only version of STRAND, which is on par with ProRNA3D-single for predictions below 10Å and slightly worse for the percentage of samples below a cRMSD of 5Å. For ProRNA3D-single, we also observe the strongest improvements, with decreased cRMSD scores for 84%, 77.7%, and 84% of the targets for STRAND_tr_, STRAND_rot_, and STRAND_tr+rot_, respectively. Please find visualizations for individual modeling tasks in Appendix C.3.

**Table 2.** RNA-protein backbone refinement results of STRAND for ProRNA3D-single predictions on PDB structures validated by **X-ray** crystallography. We show results for the translation only model STRAND_tr_, the rotation only model STRAND_rot_, the torsion only model STRAND_tor_, and the combined versions STRAND_tr+rot_ and STRAND_tr+rot+tor_. Results are reported in terms of mean and median complex RMSD (cRMSD; lower is better) and percentage of samples below a certain cRMSD threshold (2Å, 5Å, 10Å ; higher is better). We observe that all methods improve the prediction quality of ProRNA3D-single on average, with the largest improvement observed by STRAND_tr+rot_. Remarkably, the models trained with torsion can improve the mean cRMSD of the ProRNA3D-single predictions.

| Methods | Complex RMSD (Å) ↓ |  |  |  |  |
| --- | --- | --- | --- | --- | --- |
|  | %<2Å | %<5Å | %<10Å | Median | Mean |
| ProRNA3D-single | 4.76 | 36.51 | 57.14 | 8.09 | 12.02 |
| STRAND<sub>tr</sub> | **14.29** | **39.68** | 58.73 | 8.22 | 11.52 |
| STRAND<sub>rot</sub> | 12.70 | **39.68** | **60.32** | **6.90** | 11.30 |
| STRAND<sub>tor</sub> | 3.17 | 34.92 | 57.14 | 8.18 | 12.01 |
| STRAND<sub>tr+rot</sub> | 12.70 | 38.10 | 58.73 | 7.25 | **11.19** |
| STRAND<sub>tr+rot+tor</sub> | 7.94 | 38.10 | 58.73 | 7.70 | 11.49 |

For our subsequent experiments and analysis we decide to use the translation only model STRAND_tr_, the rotation only model STRAND_rot_ and their combination STRAND_tr+rot_ as these showed remarkable refinement capabilities for both AlphaFold 3 and ProRNA3D-single.

### 4.2. Sample Selection for STRAND Predictions

In the previous section, we observe remarkable refinement capabilities of STRAND, improving AlphaFold 3 and ProRNA3D-single predictions across all datasets and metrics. However, the sample selection in our initial study was done manually based on the cRMSD across 40 samples drawn from the individual STRAND models. Selecting the best sample from a set of predictions is a challenging task. We decide to follow Ketata et al. (2023) and train a confidence model based on the predictions of STRAND_tr+rot_. However, in contrast to Ketata et al. (2023), we do not train a classifier to select samples below a certain ligand RMSD threshold (5Å for Ketata et al. (2023)) but directly train a regression model for the prediction of ligand RMSDs. In addition, we cannot train our confidence model at the same scale as Ketata et al. (2023) due to compute limitations; we only use 3% of the training samples compared to Ketata et al. (2023) and train the confidence model for 60 hours on a single A40 GPU. One advantage of STRAND is that we can always fall back to the initial structure predictions of AlphaFold 3 or ProRNA3D-single in case STRAND cannot improve the prediction, since we are only doing refinement. Therefore, we include the initial predictions in the set of samples of the different STRAND versions, essentially ranking 41 instead of 40 samples per complex prediction task with the confidence model. Following Corso et al. (2022), we evaluate the confidence model in terms of top1, top5, as well as additional top10, and top20 performance, where topN refers to selecting the most accurate pose out of the N highest ranked predictions by the confidence model. In the following, we discuss the sample selection results for the three best models STRAND_tr_, STRAND_rot_, and STRAND_tr+rot_.

#### 4.2.1. Selection forAlphaFold3 Refinements

We show the results for the sample selection for STRAND using the confidence model for the refined predictions of AlphaFold 3 on non-X-ray data in Table 3. The results for X-ray data can be found in Table 10 in Appendix C.1. Generally, we observe that the confidence model can select strong samples well for the non-X-ray dataset (see Table 3). STRAND_tr_ already improves the AlphaFold 3 median performance for the top1 selected sample (6.81Å and 7.15Å for STRAND_tr_ and AlphaFold 3, respectively), while both other STRAND versions perform equally well or only slightly worse in terms of median cRMSD. From top5 onwards, all STRAND versions consistently show lower median cRMSDs, while STRAND_tr_ and STRAND_rot_ also outperform AlphaFold 3 in terms of mean performance. However, sample selection on the X-ray dataset appears more challenging. None of the STRAND versions can improve AlphaFold 3 performance for top1 and top5 selected samples, while STRAND_rot_ is the only model that can improve AlphaFold 3 mean performance for top10 selection with equal median cRMSD. However, for top20 selection, we still observe improvements with STRAND across all models for mean and median cRMSDs, except for STRAND_tr_ with slightly higher mean cRMSD but the lowest median cRMSD.

**Table 3.** RNA-protein backbone refinement results of STRAND for AlphaFold 3 **non-X-ray** predictions using sample selection with a trained confidence model. We show results for STRAND_tr_, STRAND_rot_ and the combined version STRAND_tr+rot_. Best performance is indicated in bold, equal best performance is indicated by underlined scores. Results are reported in terms of mean and median cRMSD (lower is better) after sample selection with the confidence model for STRAND compared to the cRMSD achieved by the initial AlphaFold 3 prediction.

| Methods | Complex RMSD (Å) ↓ |  |  |  |  |  |  |  |
| --- | --- | --- | --- | --- | --- | --- | --- | --- |
|  | Top 1 |  | Top 5 |  | Top 10 |  | Top 20 |  |
|  | Median | Mean | Median | Mean | Median | Mean | Median | Mean |
| AlphaFold 3 | 7.15 | **10.86** | – | – | – | – | – | – |
| STRAND<sub>tr</sub> | **6.81** | 11.59 | **6.78** | 10.75 | **6.78** | 10.67 | **6.58** | 10.58 |
| STRAND<sub>rot</sub> | 7.24 | 11.56 | 7.03 | **10.73** | 7.03 | **10.51** | 6.80 | **10.27** |
| STRAND<sub>tr+rot</sub> | 7.15 | 12.81 | 6.95 | 11.15 | 6.95 | 11.94 | 6.94 | 10.48 |

We attribute the difference in sample selection performance between the non-X-ray and the X-ray dataset to the overall high prediction quality of AlphaFold 3 and the STRAND refinements across all samples of the X-ray dataset. Distinguishing very good from good predictions seems to be much harder than selection for the non-X-ray examples, where we observed higher cRMSDs for the initial AlphaFold 3 predictions in general.

#### 4.2.2. Selection forProRNA3D-singleRefinements

The results for the sample selection for STRAND refinements of ProRNA3D-single predictions are shown in Table 4. Generally, both STRAND_tr+rot_ and STRAND_rot_ achieve similar performance, with STRAND_tr+rot_ performing best, achieving the lowest mean and median cRMSDs from top5 selection onwards (7.11Å and 8.09Å median, 11.74Å and 12.02Å mean cRMSD for STRAND_tr+rot_ top5 selection and ProRNA3D-single, respectively), while being only slightly worse than ProRNA3D-single for the top1 selected sample. While STRAND_tr_ can similarly reduce the mean cRMSD compared to ProRNA3D-single from top5 selection onwards, it cannot achieve a lower median cRMSD.

**Table 4.**
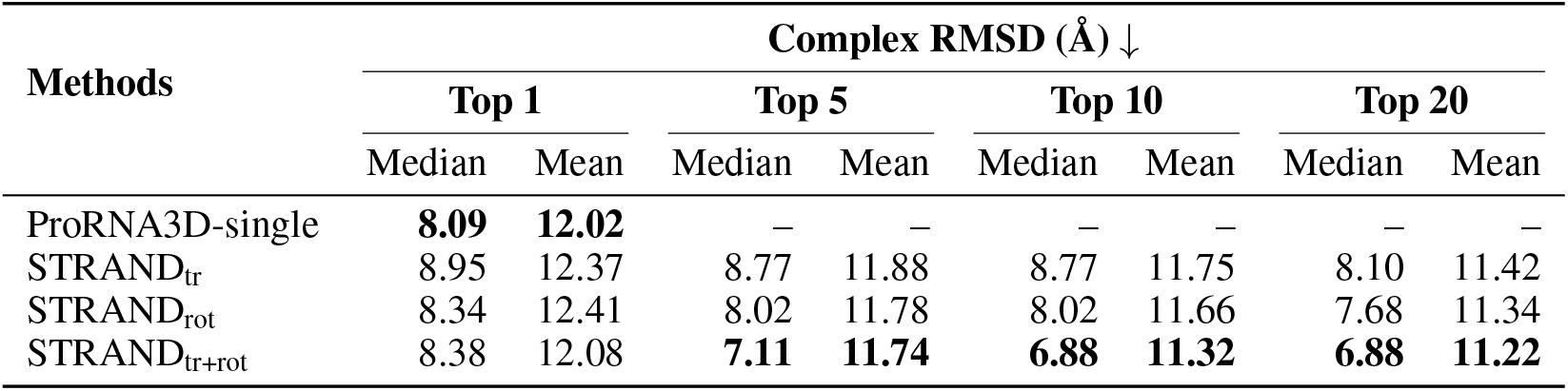
RNA-protein backbone refinement results of STRAND for ProRNA3D-single predictions for **X-ray** data using sample selection with a trained confidence model. We show results for STRAND_tr_, STRAND_rot_ and the combined version STRAND_tr+rot_. Best performance is indicated in bold. Results are reported in terms of mean and median cRMSD (lower is better) after sample selection with the confidence model for STRAND compared to the cRMSD achieved by the initial ProRNA3D-single prediction.

| Methods | Complex RMSD (Å) ↓ |  |  |  |  |  |  |  |
| --- | --- | --- | --- | --- | --- | --- | --- | --- |
|  | Top 1 |  | Top 5 |  | Top 10 |  | Top 20 |  |
|  | Median | Mean | Median | Mean | Median | Mean | Median | Mean |
| ProRNA3D-single | **8.09** | **12.02** | – | – | – | – | – | – |
| STRAND<sub>tr</sub> | 8.95 | 12.37 | 8.77 | 11.88 | 8.77 | 11.75 | 8.10 | 11.42 |
| STRAND<sub>rot</sub> | 8.34 | 12.41 | 8.02 | 11.78 | 8.02 | 11.66 | 7.68 | 11.34 |
| STRAND<sub>tr+rot</sub> | 8.38 | 12.08 | **7.11** | **11.74** | **6.88** | **11.32** | **6.88** | **11.22** |

**Table 5.** Hyperparameters of the Tensor Product Score Model.

| Parameter | Value |
| --- | --- |
| Max Ligand Radius | 5 |
| Receptor Max Radius | 30 |
| Cross Max Distance | 80 |
| Dynamic Max Cross | True |
| Cross Cutoff Weight | 3 |
| Cross Cutoff Bias | 40 |
| Center Max Distance | 30 |
| Spherical Harmonics (lmax) | 2 |
| Node Features (ns) | 16 |
| Vector Features (nv) | 4 |
| Scale by Sigma | True |
| Number of Conv Layers | 4 |
| Dropout Rate | 0 |
| Batch Normalization | False |
| Edge Features | 4 |
| Distance Embedding Dim | 32 |
| Cross Distance Embedding Dim | 32 |
| Sigma Embedding Dim | 32 |
| LM Embedding Dim | 640 |
| Hidden Features | 3 * ns |
| Gaussian Smearing Start | 0.0 |
| Gaussian Smearing Stop | 5.0 |
| Sinusoidal Max Positions | 1e4 |
| Embedding Scale | 10000 |

**Table 6.** Other Hyperparameters.

| Parameter | Value |
| --- | --- |
| Batch Size STRAND<sub>tr</sub> | 4 |
| Batch Size STRAND<sub>rot</sub> | 24 |
| Batch Size STRAND<sub>tor</sub> | 12 |
| Batch Size STRAND<sub>tr+rot</sub> | 12 |
| Batch Size STRAND<sub>tr+rot+tor</sub> | 12 |
| Number of Nearest Neighbors | 30 |
| Optimizer | Adam |

**Table 7.** RNA and Protein Residue Statistics for Different Evaluation Sets.

|  | **X-ray** | **Non-X-ray** | **ProRNA3D-single** |
| --- | --- | --- | --- |
| Num samples | 35 | 46 | 63 |
| **RNA** |  |  |  |
| Max Length | 84.0 | 828.0 | 270.0 |
| Min Length | 3.0 | 5.0 | 3.0 |
| Median Length | 6.0 | 25.5 | 10.0 |
| Mean length | 12.9 | 96.0 | 35.0 |
| **Protein** |  |  |  |
| Max length | 816.0 | 1017.0 | 850.0 |
| Min Length | 75.0 | 100.0 | 82.0 |
| Median Length | 167.0 | 669.5 | 388.0 |
| Mean Length | 289.7 | 572.3 | 416.3 |

**Table 8.** RNA and Protein Residue Statistics for Train, Validation, and Test Sets (all three combined).

|  | **Train** | **Val** | **Test** |
| --- | --- | --- | --- |
| **RNA** |  |  |  |
| Max Length | 829.0 | 706 | 240 |
| Min Length | 1.0 | 2 | 14 |
| Median Length | 22.0 | 21.0 | 68.0 |
| Mean Length | 59.0 | 60.88 | 79.0 |
| **Protein** |  |  |  |
| Max Length | 1022 | 1007 | 999 |
| Min Length | 14 | 61 | 75 |
| Median Length | 411.0 | 416.0 | 374.0 |
| Mean Length | 449.5 | 484.0 | 423.6 |

**Table 9.** RNA and Protein Residue Statistics for Train, Validation, and Test Sets used in STRAND_tr+rot+tor_.

|  | **Train** | **Val** | **Test** |
| --- | --- | --- | --- |
| **RNA** |  |  |  |
| Max Length | 1012 | 924 | 950 |
| Min Length | 2.0 | 2 | 2 |
| Median Length | 118 | 119.5 | 114.5 |
| Mean Length | 150.86 | 148.76 | 118.68 |
| **Protein** |  |  |  |
| Max Length | 1015 | 823 | 476 |
| Min Length | 2 | 50 | 51 |
| Median Length | 155 | 126.5 | 152.5 |
| Mean Length | 449.65 | 196.26 | 178.32 |

**Table 10.** RNA-protein backbone refinement results of STRAND for AlphaFold 3 **X-ray** predictions using sample selection with a trained confidence model. We show results for STRAND_tr_, STRAND_rot_ and the combined version STRAND_tr+rot_. Best performance is indicated in bold, equal best performance is indicated by underlined scores. Results are reported in terms of mean and median cRMSD (lower is better) after sample selection with the confidence model for STRAND compared to the cRMSD achieved by the initial AlphaFold 3 prediction.

| Methods | Complex RMSD (Å) ↓ |  |  |  |  |  |  |  |
| --- | --- | --- | --- | --- | --- | --- | --- | --- |
|  | Top 1 |  | Top 5 |  | Top 10 |  | Top 20 |  |
|  | Median | Mean | Median | Mean | Median | Mean | Median | Mean |
| AlphaFold 3 | **2.20** | **3.09** | — | — | — | — | — | — |
| STRAND<sub>tr</sub> | 2.78 | 3.94 | 2.39 | 3.57 | 2.30 | 3.47 | **2.11** | 3.15 |
| STRAND<sub>rot</sub> | 3.64 | 4.46 | 2.22 | 3.30 | <u>2.20</u> | **2.90** | <u>2.20</u> | **2.83** |
| STRAND<sub>tr+rot</sub> | 2.84 | 4.48 | 2.45 | 3.60 | 2.29 | 3.34 | 2.12 | 2.90 |

## 5. Discussion, Limitations & Future Work

In this work, we introduce Structure refinement of RNA-protein complexes via diffusion (STRAND), a novel approach for monomeric RNA-protein complex refinement. We leverage the capabilities of current deep learning approaches for structural biology that can model different molecule types and complexes of these and combine it with the powerful diffusion framework of DiffDock-PP. Our new method successfully refines monomeric RNA-protein complex predictions by enhancing docking positions, showing remarkable improvements in structure quality when applied to AlphaFold 3 and ProRNA3D-single predictions.

With our analysis using different individual transformations and their combinations, we show that simple rotation and translation of the initial predictions can often already lead to improvements, indicating that the 3D structure predictors might often generally misplace the protein with respect to the RNA when modeling their interaction. This is further supported by our finding that the local refinement via torsion angles does not lead to substantial improvements, although torsion angle refinement generally comes with its own additional challenges and the training of the torsion models might require further development, specifically regarding noise schedules. However, we think that our results provide new insights that could help to develop stronger 3D RNA-protein structure prediction tools in the future, which would also be beneficial for new refinement methods.

Our experiments for sample selection show that it is generally possible to select strong samples by training a confidence model for ligand RMSD prediction. We think that direct prediction of ligand RMSD could be beneficial over training a classifier to select samples below a certain ligand RMSD threshold, as implemented in DiffDock-PP. However, we follow Ketata et al. (2023) and optimize the ligand RMSD, which might not be optimal. It would be interesting to analyze the results when optimizing e.g. for cRMSD directly, since cRMSD might add information to guide the model predictions to reduce global misplacement. Nevertheless, our selection model already shows remarkable results, although we train only on a fraction of the data used by Ketata et al. (2023) due to compute limitations.

### Limitations

However, despite notable improvements in performance, our approach also has limitations. Our refinement currently only involves backbone atoms similar to DiffDock-PP. However, it would be generally preferable to refine full-atom structures. Additionally, we are bound by the length limitations of the foundation models for obtaining sequence embeddings. We currently limit the length to 1022 residues. Furthermore, selecting the best sample remains challenging. We note, however, that our confidence model is currently trained on a relatively small set of complexes, and we expect improved performance when scaling the training to larger amounts of data.

### Future Work

For the future, we mainly plan to address the current limitations. Most importantly, we would like to further analyze the inclusion of torsion angle refinement and scale the data for training the confidence model for sample selection. Also, optimizing the confidence model for cRMSD instead of ligand RMSD would be a natural next step.

We believe that our novel refinement approach bears large potential for the community, potentially enabling high prediction quality of monomeric RNA-protein complexes in the future.

## Acknowledgments

The authors acknowledge funding by the German Research Foundation (DFG) under SFB 1597 (SmallData), grant no. 499552394, and through grant no. 417962828 as well as support by the state of Baden-Württemberg through bwHPC and the German Research Foundation (DFG) through grant no INST 39/963-1 FUGG (bwForCluster NEMO) and grant INST 35/1597-1 FUGG (bwForCluster Helix). This research was funded by the European Union (via ERC Consolidator Grant DeepLearning 2.0, grant no. 101045765). Views and opinions expressed are however those of the author(s) only and do not necessarily reflect those of the European Union or the European Research Council. Neither the European Union nor the granting authority can be held responsible for them.

---

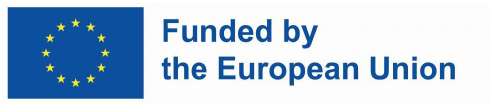

## A. Hyperparameters

## B. Data Insights

### B.1. Statistics of the three sets used in the evaluation of structure refinement

### B.2. Statistics train, validation, and test splits used in training and evaluating the diffusion model

## C. Additional Results

### C.1. Improving AlphaFold 3 Prediction for X-Ray Structures

### C.1. Improving AlphaFold 3 Prediction for Non-X-Ray Structures

### C.3. Improving ProRNA3D-single Predictions

